# Resveratrol shortens the chronological lifespan of *Saccharomyces cerevisiae* by a pro-oxidant mechanism

**DOI:** 10.1101/2020.07.17.209411

**Authors:** Juan Carlos Canedo-Santos, Iridian Mora-Martinez, Ingrid Karina Gutierrez-Garcia, Maria Guadalupe Ramirez-Romero, Luis Alberto Madrigal-Perez

## Abstract

Resveratrol consumption has linked with normalization of the risk factors of some diseases such as colon cancer and type 2 diabetes. The antioxidant phenotype caused by resveratrol has recognized as a key piece in the health benefits exerted by this phytochemical. Although the antioxidant activity showed by resveratrol has attributed at the molecule *per se*, recent evidence indicates that the antioxidant effect occasioned by resveratrol could be associated with a pro-oxidant mechanism. The hypothesis that resveratrol inhibits complex III of the electron transport chain as its main target suggests that resveratrol increases reactive oxygen species (ROS) generation produces via reverse electron transport. This idea also explains that cells respond to the oxidative damage caused by resveratrol, inducing their antioxidant systems. The free radical theory of aging postulates that organisms age due to the accumulation of the harmful effects of ROS in cells. For these reasons, we hypothesize that resveratrol shortens the chronological life span (CLS) of *Saccharomyces cerevisiae* due to a pro-oxidant activity. Herein, we provide evidence that 100 μM resveratrol supplementation at 5% glucose: 1) shorted the CLS of *CTT1* and *YAP1* genes deleted strains; 2) decreased the H_2_O_2_ release in the WT strain, and maintain unaltered the H_2_O_2_ release in the *ctt1*Δ strain; 3) lessened exponential growth of *ctt1*Δ strain, which was reverted with the adding of GSH; 4) increased catalase activity in the WT strain, a phenotype that was not observed in the *ctt1*Δ strain. Altogether, these results indicate that resveratrol decreases CLS by a pro-oxidant mechanism.

## Introduction

The resveratrol (3,5,4 ′ -trihydroxy-trans-stilbene) is a polyphenol synthesized by some plants such as *Vitis vinifera* under biotic or abiotic stresses [1, 2]. Resveratrol synthesis forms part of a chemical defense mechanism of plants to counteract or prevent infections [3, 4]. Cytotoxic properties of resveratrol are well-documented and fit with its biological function in plants [5]. Another important effect exerted by resveratrol is the antioxidant, which has linked with the health benefits exerted by this compound [6]. However, the antioxidant properties of resveratrol are pro-survival to the cells and do not match with the toxic effect shown by resveratrol. Even the antioxidant molecular mechanism of resveratrol is not yet fully understood. The hypothesis that resveratrol inhibits the oxidative phosphorylation has gained attention in the last years and explains both its toxic and the antioxidant properties [7].

The inhibition of the electron transport chain (ETC) between complex I and complex III by resveratrol impair mitochondrial respiration [8]. The inhibition of the ETC by resveratrol could maintain the ubiquinone pool and complex I more reduced, increasing the chances of electron leaking and the formation of reactive oxygen species (ROS) [7]. In support of this idea, it has been reported that 5 mg/kg of resveratrol supplementation increases lipid peroxidation of the heart, liver, and kidney of rats [9]. Additionally, thyroid carcinoma THJ-16T cell line supplemented with 100 μM resveratrol increases superoxide anion production at the mitochondrial level [10]. The murine liver connective tissue cells GRX also augmented the ROS generation when they were supplemented with 50 μM of resveratrol at 24 h and 120 h [11]. The increase of the ROS generation by resveratrol could induce the expression of the antioxidant systems as a defense mechanism to counteract this oxidative damage. In this regard, it has reported that *Saccharomyces cerevisiae* cells exposed to 5 μM of resveratrol accumulated ROS and deletion of the antioxidant transcription factor gene *YAP1* in these yeast cells increased cellular toxicity of resveratrol [12]. Also, in primary epidermal keratinocytes derived from human skin, 50 μM of resveratrol supplementation triggered antioxidant systems transcription, dependent on the transcription factor Nrf2 [13]. Interestingly, resveratrol also augmented superoxide anion generation and intracellular ROS production in keratinocytes cells [13]. In accordance with the free radical theory of aging [14], resveratrol supplementation could shorten the chronological life span (CLS) of *S. cerevisiae* by a pro-oxidant mechanism.

Resveratrol also exerts phenotypes in a dose-dependent manner, fitting well with the hormetic behavior, as has been reported for cellular viability [15] and pro-oxidative properties [13]. Importantly, a diet-dependent effect of phenotypes exerted by resveratrol has also been documented [5]. For example, resveratrol exhibit a glucose-dependent effect in chronological aging [8], cellular viability [16], mitochondrial respiration [16], and hydrogen peroxide release [8] in *S. cerevisiae.* In the Apc^Min^ mice, a model of colorectal carcinogenesis, resveratrol treatment (0.00007%) decrease adenoma number per mouse in a high-fat diet but not in a standard diet [17]. However, the evidence linking the pro-oxidant properties of resveratrol with a glucose-dependent mechanism is still lacking.

For these reasons, this study aimed to demonstrate that resveratrol causes a pro-oxidant response that impact in chronological longevity, cell growth, and ROS generation in a glucose concentration-dependent manner. Herein, we provide evidence that resveratrol supplementation shortens CLS of *ctt1*Δ and *yap1*Δ strains at 5% glucose. None effect was observed in CLS at 0.5% of glucose in *ctt1*Δ*, yap1*Δ*, hcm1*Δ, *sod2*Δ, and *msn2*Δ deletant strains, with resveratrol supplementation. The deletion of the *CTT1* gene reverts the decrease in the H_2_O_2_ release and the increase in catalase activity occasioned by resveratrol. At 5% glucose, resveratrol supplementation diminishes the growth of the *ctt1*Δ strain, and this phenotype was reestablished with the adding of reduced glutathione.

## Material and methods

### Strains

To perform the experiments was used *S. cerevisiae* BY4742 strain (*MAT*α; *his3Δ1; leu2Δ0; lys2Δ0; ura3Δ0*) and its deletant strains in the genes *YAP1* (*MAT*α*; his3Δ1; leu2Δ0; lys2Δ0; ura3Δ0; YML007w::kanMX4*), *CTT1* (*MAT*α*; his3Δ1; leu2Δ0; lys2Δ0; ura3Δ0; YGR088w::kanMX4*), *MSN2* (*MAT*α*; his3Δ1; leu2Δ0; lys2Δ0; ura3Δ0; YMR037c::kanMX4*), *SOD2*(*MAT*α*; his3Δ1; leu2Δ0; lys2Δ0; ura3Δ0; YHR008c::kanMX4*) and *HCM1* (*MAT*α*; his3Δ1; leu2Δ0; lys2Δ0; ura3Δ0; YCR065w::kanMX4*) obtained from EURSOCARF. The strains were maintained in yeast extract-peptone-dextrose (YPD) medium (1% yeast extract, 2% casein peptone and 2% glucose), deletant strains were supplemented with geneticin (G-418 disulfate salt solution, Sigma-Aldrich, St. Louis, MO, USA) at a final concentration of 200 μg/mL.

### Chronological lifespan assay

The CLS was determined according to Ramos-Gomez, Olivares-Marin, Canizal-Garcia, Gonzalez-Hernandez, Nava and Madrigal-Perez [8]. Briefly, the synthetic-complete (SC) medium was used to perform the CLS assay. It consisted of 0.18% yeast nitrogen base without amino acids, 0.5% ammonium sulfate, 0.2% KH_2_PO_4_, 1% drop-out mix without uracil, supplemented with 400 μg/mL of uracil. The mediums were supplemented with two different glucose concentrations (0.5% and 5%) and five levels of resveratrol (0, 0.1, 1, 10, 100, and 1000 μM; *trans*-resveratrol ≥ 99% HPLC, Sigma-Aldrich) that were added at the beginning of the CLS assay. Afterward, 1 mL of the SC medium was inoculated with 1% of a fresh overnight culture of *S. cerevisiae* in a 15 mL conic tube. The *S. cerevisiae* cultures were grown at 30 °C with constant shaking at 250 rpm for 15 days. Once three days of incubation were passed, aliquots of 5 μL were taken and inoculated into 145 μL YPD 2% glucose, every two days. Samples were placed in a 96-well plate and incubated at 30 °C for 24 h in a Varioskan Sky (Thermo-Scientific, Waltham, MA, USA) programmed with continuous shaking and readings at 600 nm each 60 min. The survival percentage (*Sn*) was calculated according to equation 1:

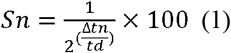

Where *Δtn* is the time shift (h), obtained by interpolation of D.O._600_=0.5 with a linear regression of the exponential phase of the growth curve, and *td* is the doubling time (h).

### Growth kinetics

The cell growth was calculated using exponential growth as an indirect marker of cell division, as described by Olivares-Marin, Madrigal-Perez, Canizal-Garcia, Garcia-Almendarez, Gonzalez-Hernandez and Regalado-Gonzalez [18]. Growth kinetics were begun at O.D._600_ •0.1 in a 25 mL shake flasks that contained 5 mL of YPD media supplemented with 5% glucose and 100 μM resveratrol. The shake flasks were incubated in a MaxQ 4000 incubated shaker (Thermo-Scientific) at 30 °C for 12 hours with shaking at 250 rpm. The growth was monitored, measuring the O.D._600_ each hour. Data were analyzed using the statistical package GraphPad Prism 6.00 for Macintosh (GraphPad Software), fitting the growth kinetics with the exponential growth equation to obtain the specific growth rate (*μ*).

### Hydrogen peroxide release

Hydrogen peroxide (H_2_O_2_) release was quantified as an indicator of ROS production using the Amplex red hydrogen peroxide assay kit (Invitrogen, Waltham, MA, USA) following the manufacturer instructions. Briefly, exponential-growth phase *S. cerevisiae* cultures (O.D._600_ ~ 0.6) grown in SC medium with 5% glucose were harvested at 5000 x *g* for 5 minutes at 28 °C and washed two times with 5 mL of sterile deionized water at 28 °C. Then, cellular pellets were resuspended in 2 mL of assay buffer containing 20 mM Tris-HCl, 0.5 mM EDTA, 2% of ethanol at pH 7. Finally, cells were placed into a 96-well plate at a density of 3 × 10^6^ cells/well. The microplate was incubated at 30 °C with constant agitation for 30 min. The basal release of H_2_O_2_ was measured at a wavelength of 560 nm with a microplate reader (Varioskan Sky, Thermo-Scientific).

### Catalase activity

Protein isolation for catalase activity assay was performed in exponential-growth phase *S. cerevisiae* cultures (O.D._600_ ~ 0.6) grown in SC medium supplemented with 5% glucose and 100 μM of resveratrol. Cultures were harvest at 13000 x *g* for 3 minutes at 4 °C and resuspended in 500 μL of sterile distilled water. Then, it was added 500 μL of 0.7 N sodium hydroxide and incubated for 5 minutes at room temperature. Afterward, 0.2 g of glass beads (0.2 mm) were placed into the tubes and vortexing 1 minute and cooled into ice 2 minutes, repeating this two times. The solution was centrifuged at 5000 x *g* for 1 minute, discarding the supernatant. The pellet was resuspended in a solution consisted of 100 mM of potassium sodium, 5 mM of EDTA, and 1 mM of 2-mercaptoethanol. Next, the solution was centrifuged at 3500 x *g* for 1 min; the supernatant was used as the total protein isolation. For quantify catalase activity, 2 μL of the total protein isolation was mixed with 0.5 μL of 0.2 M H_2_O_2_ and placed into μDrop plate. The plate was incubated at 25 °C for 5 minutes in a Varioskan Sky microplate reader (Thermo-Scientific), recording the absorbance at 240 nm at the initial and final of the incubation period. The catalase activity was normalized with the total protein quantification.

### Statistical analyses

The mean ± standard deviation from at least three independent experiments was graphed. Means were compared using one-way ANOVA followed by a Dunnett multiple comparisons to analyze differences in the area under the curve from CLS assays and exponential growth. To analyze differences in the H_2_O_2_ release and catalase activity was used a two-tailed unpaired *t*-test. Statistical analyses were computed in the software GraphPad Prism 6.00 for Macintosh (GraphPad Software).

## Results

### Influence of resveratrol in the chronological aging of deletant strains in antioxidant systems

To evaluate whether the shorten of CLS caused by resveratrol is coming from a pro-oxidant mechanism; we decided to measure the CLS in deletant strains in genes related to the antioxidant response of *S. cerevisiae.* The experiment was conducted with two glucose concentrations: 0.5% and 5%, which promoting a respiratory and fermentative metabolism, respectively [18]. Additionally, it was used five levels of resveratrol in a logarithmic scale (0.1, 1, 10, 100, and 1000 μM) to observe a dose-dependent effect.

The deletant strain of the gene encoding the cytosolic catalase T (*CTT1*) showed a decrease in the CLS at 5% glucose when it was supplemented with 100 and 1000 μM of resveratrol in comparison with the *ctt1*Δ vehicle control (**Fig. 1i-j**). Low doses of resveratrol (0.1, 1, and 10) have did not affect the CLS at 5% glucose in the *ctt1*Δ strain (**Fig. 1f-h**). At 0.5% glucose, resveratrol did not modify the CLS when the *CTT1* gene was deleted (**Fig. 1a-e**).

**Fig 1.**
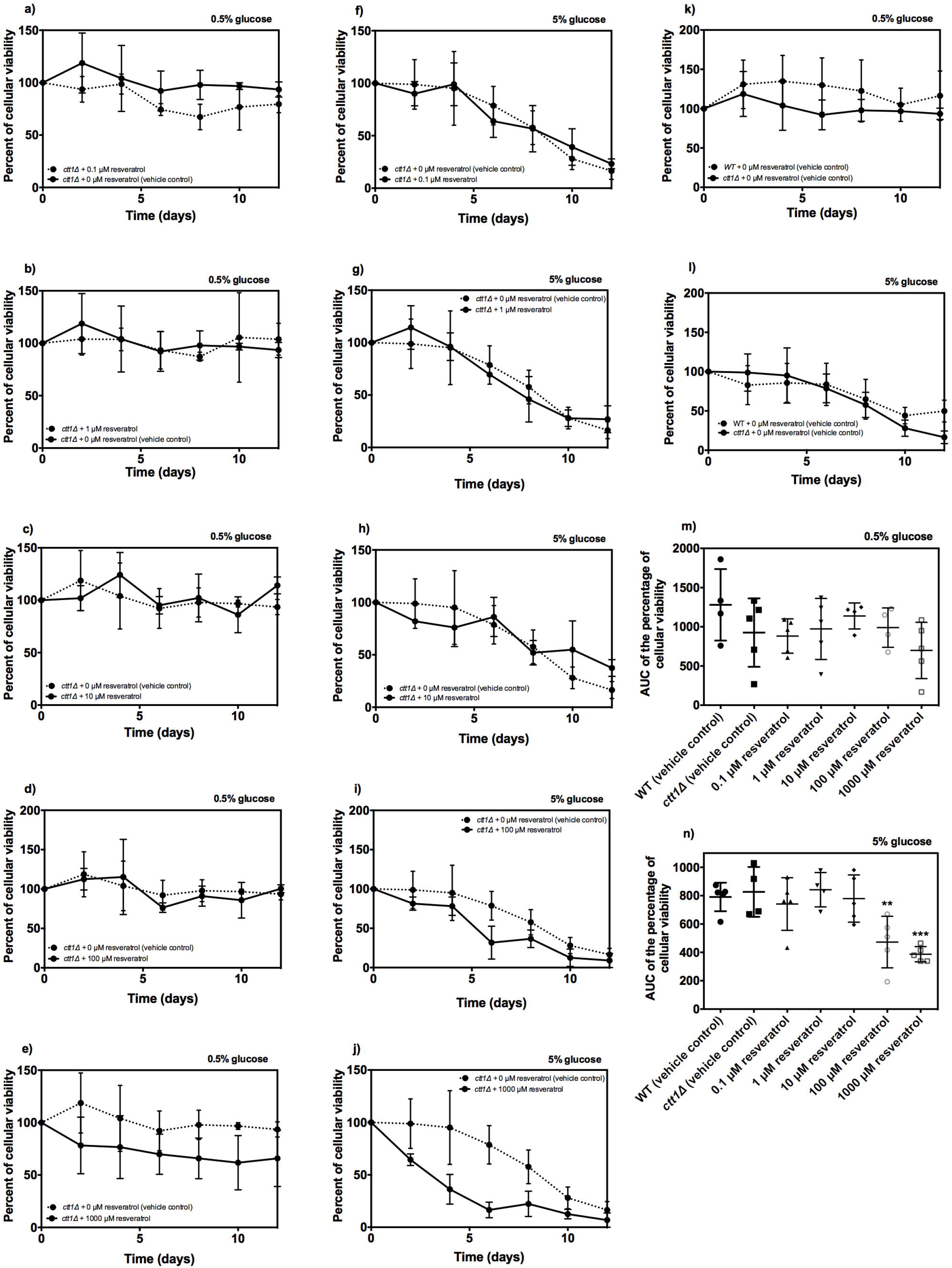
Effect of deletion of *CTT1* gene on the CLS of *S. cerevisiae* supplemented with resveratrol. The CLS was assayed in SC-medium using two glucose concentrations (0.5% and 5%) and five levels of resveratrol (0, 0.1, 1, 10, 100, and 1000 μM). **a)-e)** CLS of *ctt1*Δ strain grown with 0.5% glucose and with 0, 0.1, 1, 10, 100, and 1000 μM of resveratrol, respectively; **g)-j)** CLS of *ctt1*Δ strain grown with 5% glucose and with 0, 0.1, 1, 10, 100, and 1000 μM of resveratrol, respectively; **k)-l)** CLS comparison between *ctt1*Δ and WT strain at 0.5%, and 5% glucose, respectively; **m)-n)** Represents the area under the curve (AUC) of CLS assays at 0.5% and 5%, respectively. The AUC survival was calculated from the data of the percentage of cellular viability vs. time using the trapezoidal rule in the GraphPad Prism 6.00. The results represent mean values ± standard deviation from four to five independent experiments, which include mean values of three technical repetitions. Statistical analyses were performed using one-way ANOVA followed by Dunnett’s test *vs. ctt1*Δ (vehicle control), ** *P* ≤ 0.01; *** *P* ≤ 0.001.

The Yap1p is a basic leucine zipper transcription factor, required for the response to the oxidative stress in *S. cerevisiae* [19]. The deletion of the *YAP1* gene occasioned a decrease in the CLS of *S. cerevisiae* grown at 5% glucose, with 100 and 1000 μM of resveratrol (**Fig. 2i-j**). However, supplementation with 0.1, 1, and 10 μM of resveratrol did not modify the CLS of the *yap1*Δ strain at 5% glucose (**Fig. 2f-h**). Under the low-glucose concentration (0.5%), resveratrol supplementation did not affect the CLS of the *yap1*Δ strain (**Fig. 2a-e**).

**Fig 2.**
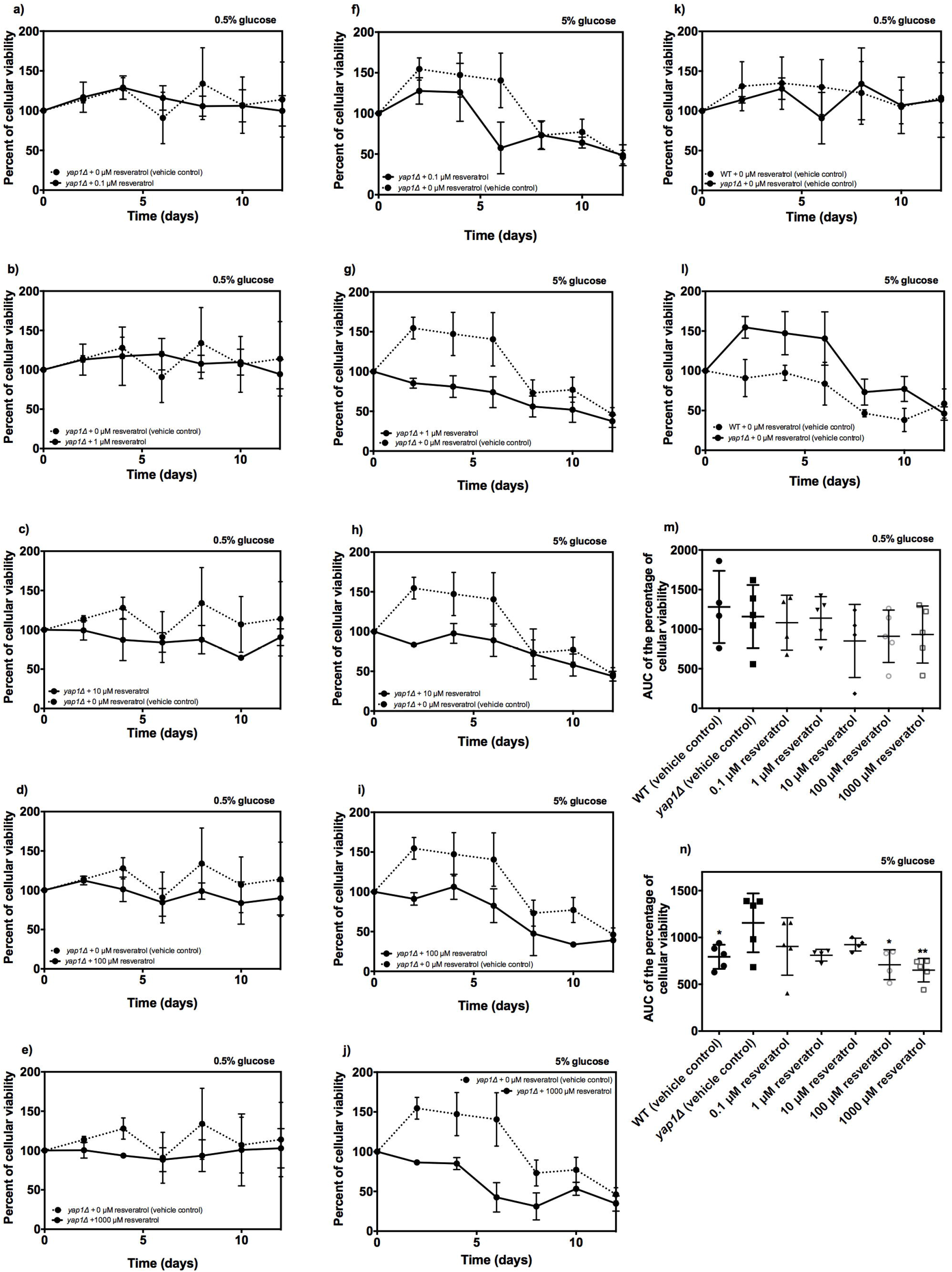
Influence of *YAP1* gene deletion on the CLS of *S. cerevisiae* supplemented with resveratrol. The CLS was assayed in SC-medium using two glucose concentrations (0.5% and 5%) and five levels of resveratrol (0, 0.1, 1, 10, 100, and 1000 μM). **a)-e)** CLS of *yap1*Δ strain grown with 0.5% glucose and with 0, 0.1, 1, 10, 100, and 1000 μM of resveratrol, respectively; **g)-j)** CLS of *yap1*Δ strain grown with 5% glucose and with 0, 0.1, 1, 10, 100, and 1000 μM of resveratrol, respectively; **k)-l)** CLS comparison between *yap1*Δ and WT strain at 0.5%, and 5% glucose, respectively; **m)-n)** Represents the area under the curve (AUC) of CLS assays at 0.5% and 5%, respectively. The AUC survival was calculated from the data of the percentage of cellular viability vs. time using the trapezoidal rule in the GraphPad Prism 6.00. The results represent mean values ± standard deviation from four to five independent experiments, which include mean values of three technical repetitions. Statistical analyses were performed using one-way ANOVA followed by Dunnett’s test *vs. yap1*Δ (vehicle control), * *P* ≤ 0.05; ** *P* ≤ 0.01.

The transcriptional factors Hcm1p and Msn2p participate in the response of *S. cerevisiae* to oxidative stress [19, 20]. Nonetheless, the deletion of the genes *HCM1* and *MSN2* did not change the CLS of *S. cerevisiae* at 0.5% or 5% glucose with any of the resveratrol concentrations tested (**Fig. 3** **and** **4**). Finally, the gene deletion of the mitochondrial manganese superoxide dismutase (*SOD2*) did not affect the CLS of *S. cerevisiae* at any glucose or resveratrol concentration (**Fig. 5**). Altogether, these results suggest that deletion *CTT1* and *YAP1* genes exacerbate the shorten of CLS caused by resveratrol supplementation.

**Fig 3.**
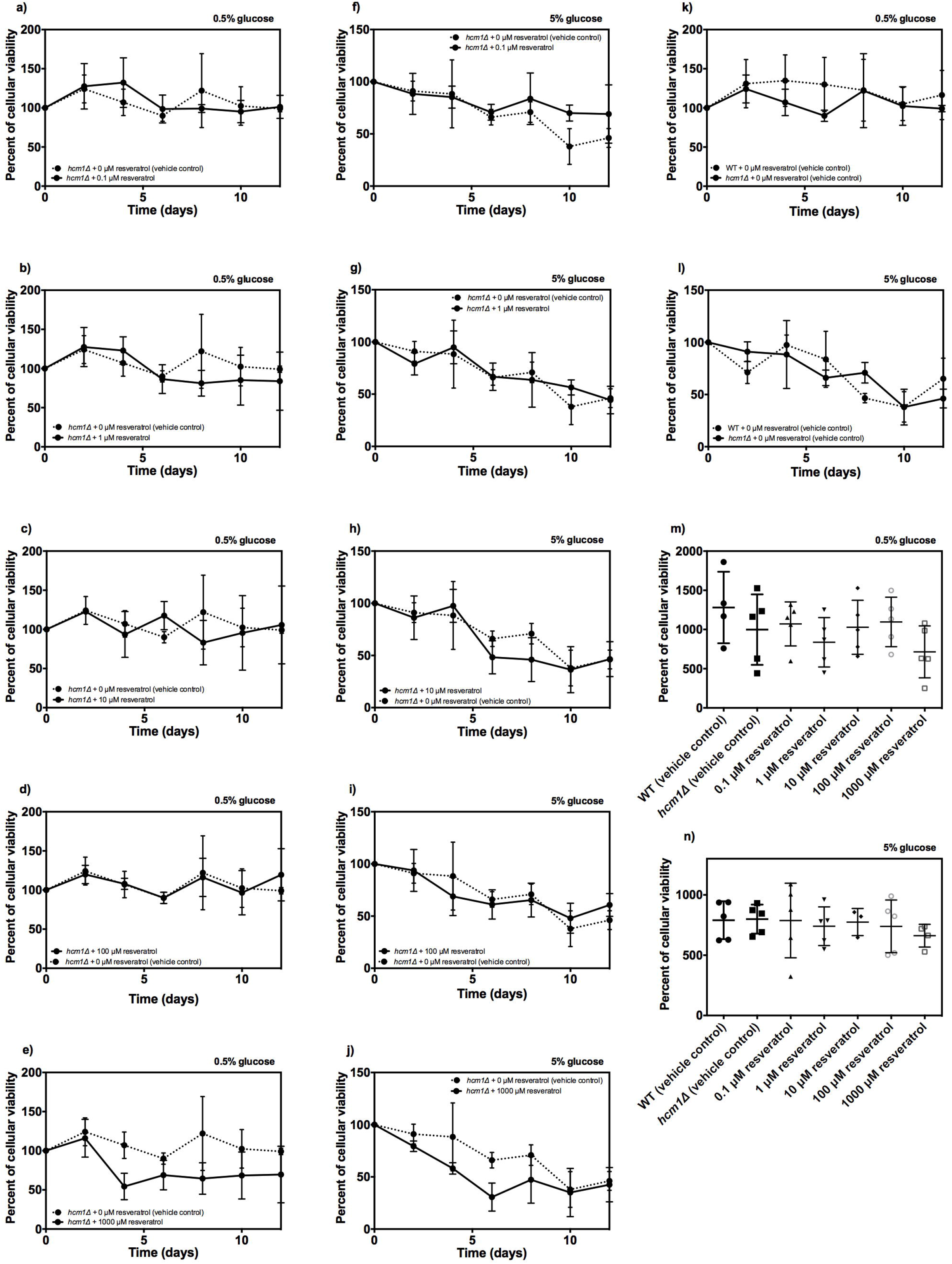
Impact of *HCM1* gene deletion on the CLS of *S. cerevisiae* supplemented with resveratrol. The CLS was assayed in SC-medium using two glucose concentrations (0.5% and 5%) and five levels of resveratrol (0, 0.1, 1, 10, 100, and 1000 μM). **a)-e)** CLS of *hcm1*Δ strain grown with 0.5% glucose and with 0, 0.1, 1, 10, 100, and 1000 μM of resveratrol, respectively; **g)-j)** CLS of *hcm1*Δ strain grown with 5% glucose and with 0, 0.1, 1, 10, 100, and 1000 μM of resveratrol, respectively; **k)-l)** CLS comparison between *hcm1*Δ and WT strain at 0.5%, and 5% glucose, respectively; **m)-n)** Represents the area under the curve (AUC) of CLS assays at 0.5% and 5%, respectively. The AUC survival was calculated from the data of the percentage of cellular viability vs. time using the trapezoidal rule in the GraphPad Prism 6.00. The results represent mean values ± standard deviation from four to five independent experiments, which include mean values of three technical repetitions. Statistical analyses were performed using one-way ANOVA followed by Dunnett’s test *vs. hcm1*Δ (vehicle control).

**Fig 4.**
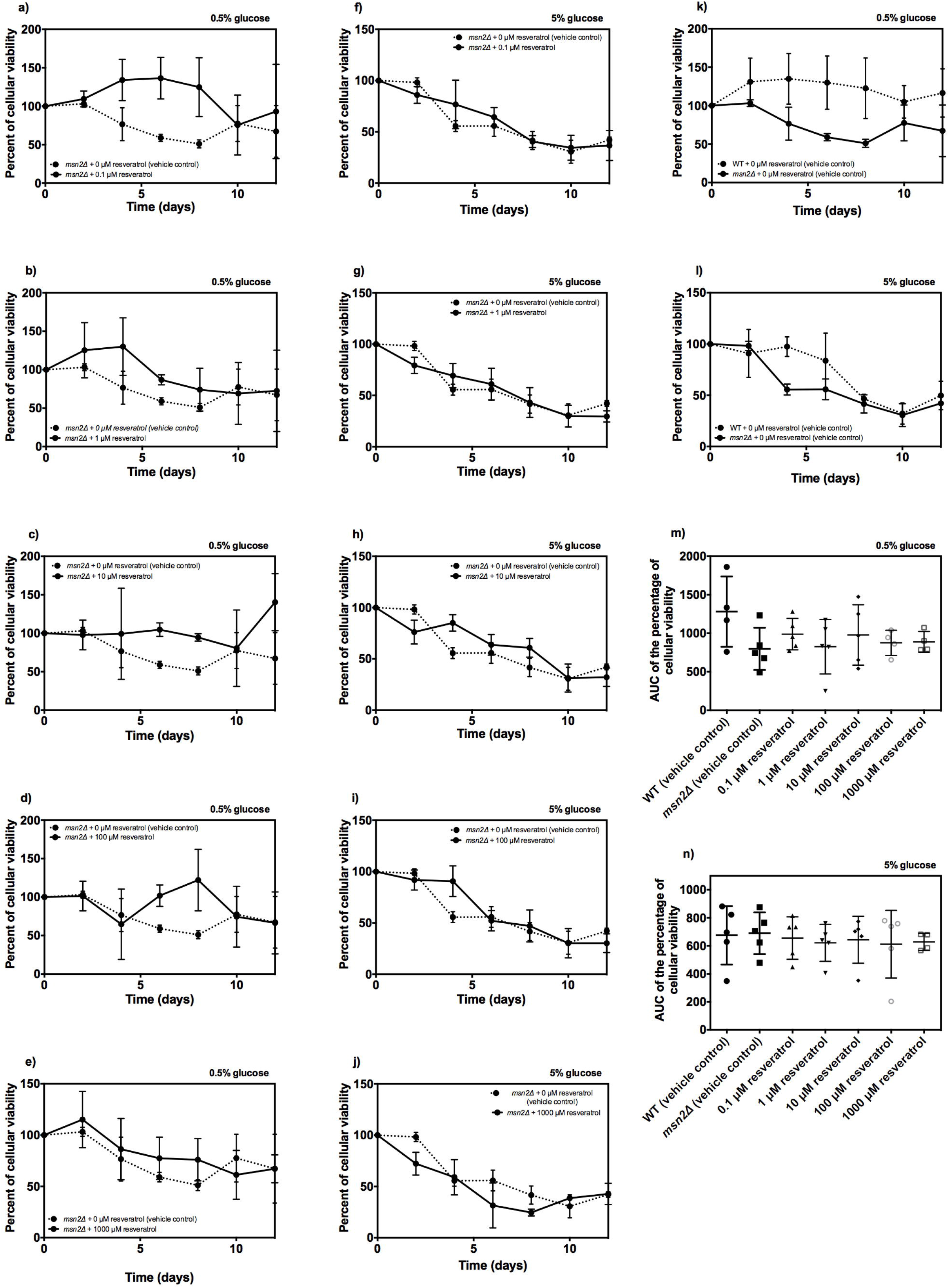
Effect of *MSN2* gene deletion on the CLS of *S. cerevisiae* supplemented with resveratrol. The CLS was assayed in SC-medium using two glucose concentrations (0.5% and 5%) and five levels of resveratrol (0, 0.1, 1, 10, 100, and 1000 μM). **a)-e)** CLS of *msn2*Δ strain grown with 0.5% glucose and with 0, 0.1, 1, 10, 100, and 1000 μM of resveratrol, respectively; **g)-j)** CLS of *msn2*Δ strain grown with 5% glucose and with 0, 0.1, 1, 10, 100, and 1000 μM of resveratrol, respectively; **k)-l)** CLS comparison between *msn2*Δ and WT strain at 0.5%, and 5% glucose, respectively; **m)-n)** Represents the area under the curve (AUC) of CLS assays at 0.5% and 5%, respectively. The AUC survival was calculated from the data of the percentage of cellular viability vs. time using the trapezoidal rule in the GraphPad Prism 6.00. The results represent mean values ± standard deviation from four to five independent experiments, which include mean values of three technical repetitions. Statistical analyses were performed using one-way ANOVA followed by Dunnett’s test *vs. msn2*Δ (vehicle control).

**Fig 5.**
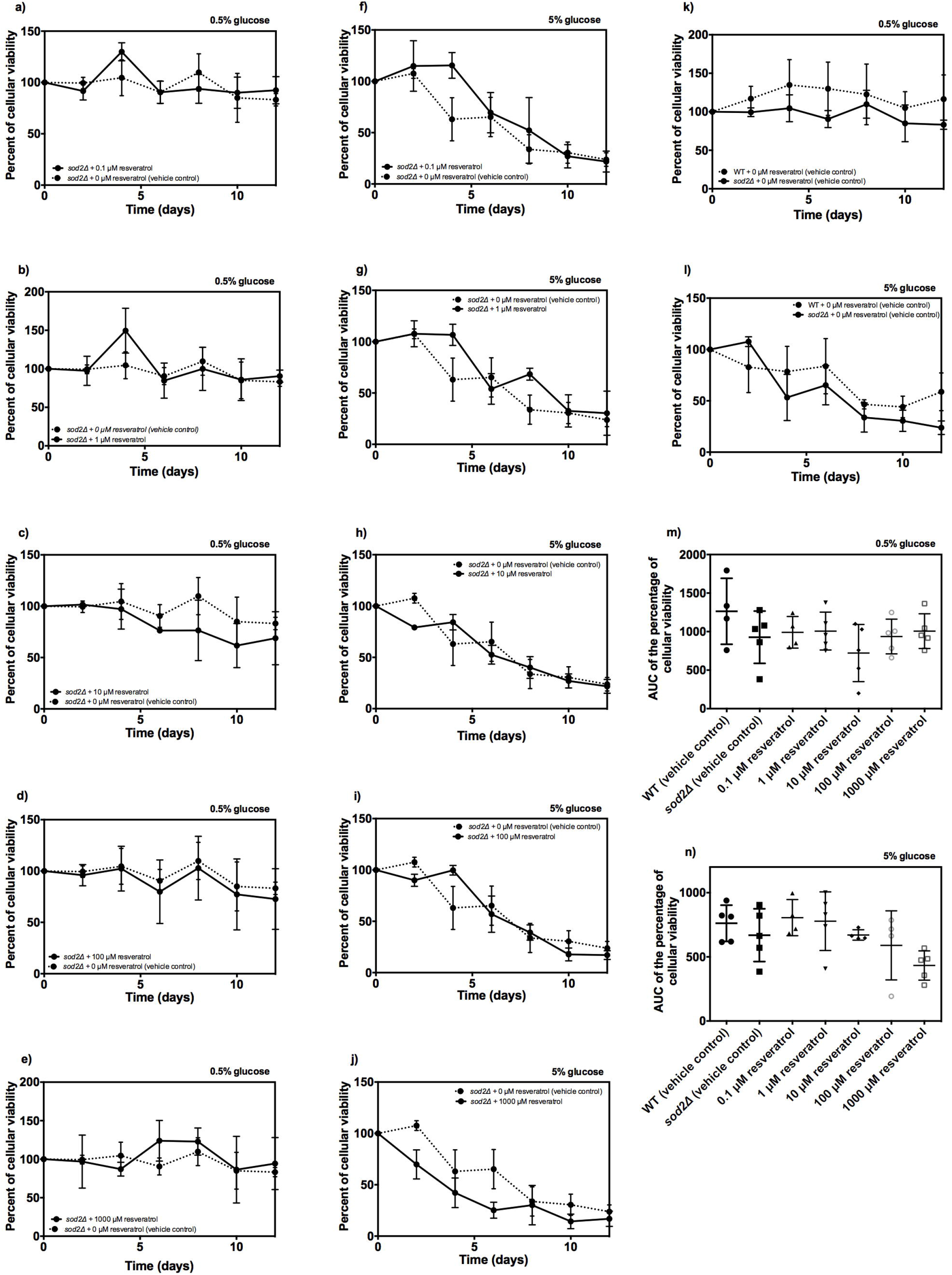
Influence of deletion *SOD2* gene on the CLS of *S. cerevisiae* supplemented with resveratrol. The CLS was assayed in SC-medium using two glucose concentrations (0.5% and 5%) and five levels of resveratrol (0, 0.1, 1, 10, 100, and 1000 μM). **a)-e)** CLS of *sod2*Δ strain grown with 0.5% glucose and with 0, 0.1, 1, 10, 100, and 1000 μM of resveratrol, respectively; **g)-j)** CLS of *sod2*Δ strain grown with 5% glucose and with 0, 0.1, 1, 10, 100, and 1000 μM of resveratrol, respectively; **k)-l)** CLS comparison between *sod2*Δ and WT strain at 0.5%, and 5% glucose, respectively; **m)-n)** Represents the area under the curve (AUC) of CLS assays at 0.5% and 5%, respectively. The AUC survival was calculated from the data of the percentage of cellular viability vs. time using the trapezoidal rule in the GraphPad Prism 6.00. The results represent mean values ± standard deviation from four to five independent experiments, which include mean values of three technical repetitions. Statistical analyses were performed using one-way ANOVA followed by Dunnett’s test *vs. sod2*Δ (vehicle control).

### H_2_O_2_ release in deletant strains in the antioxidant systems with resveratrol supplementation

To evaluate whether the decrease in CLS is related to a ROS production occasioned by resveratrol, a quantification of H_2_O_2_ release was carried out. For the following experiments, we decided to use only 100 of μM resveratrol, the lowest concentration that shortened the CLS and 5% glucose, concentration in which is observed the CLS lessening. Resveratrol supplementation decreased the H_2_O_2_ release in comparison with the vehicle control in the WT strain (**Fig. 6a**); the same phenotype was displayed in the strains *hcm1*Δ, *sod2*Δ, *yap1*Δ, and *msn2*Δ (**Fig. 6b-e**). However, the *ctt1*Δ strain did not show a difference in comparison to vehicle control in the H_2_O_2_ release at 5% glucose (**Fig. 6f**). These data suggest the cytosolic catalase T is essential to decrease the H_2_O_2_ release prompted by resveratrol supplementation, and this could be impacting the CLS.

**Fig 6.**
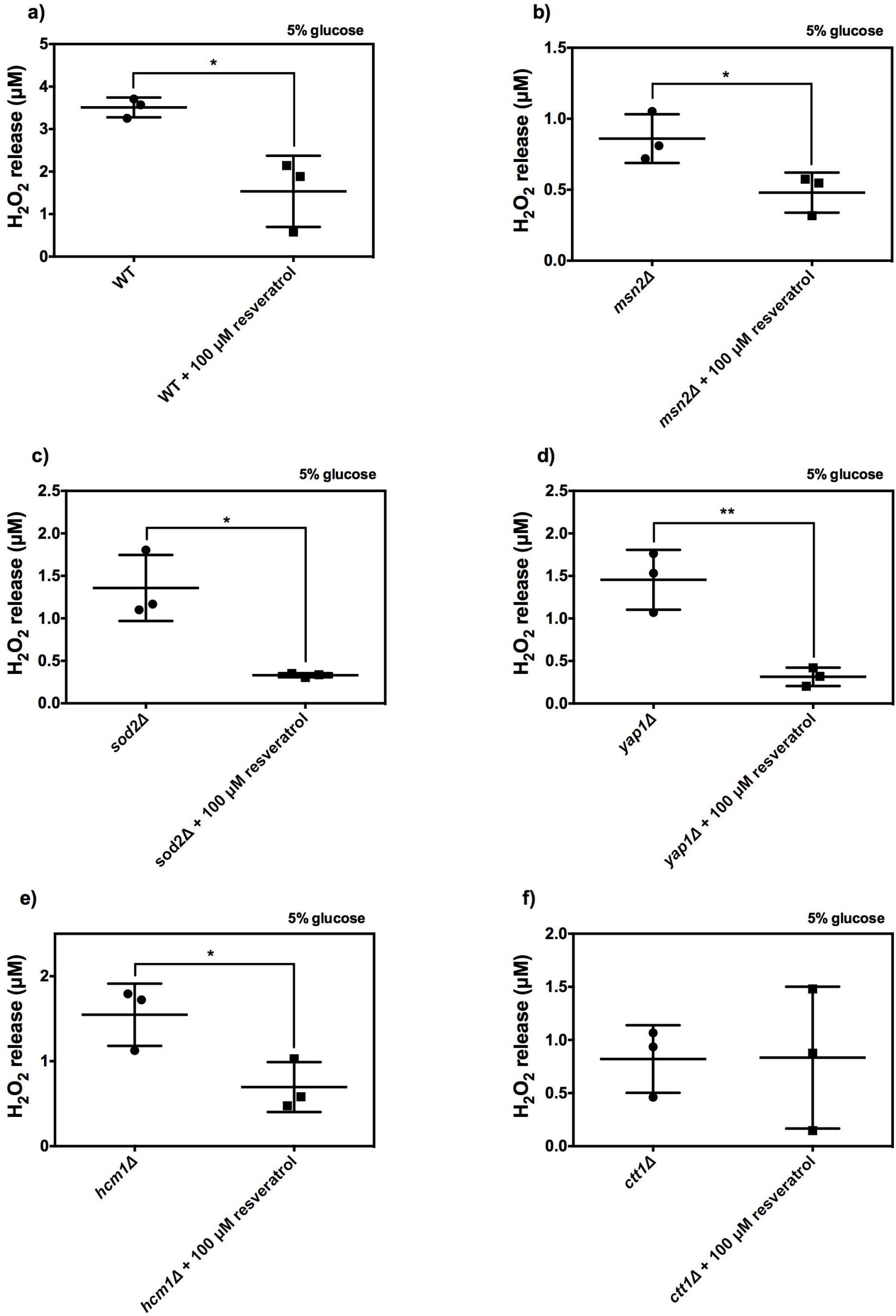
Effect of the deletion of genes *CTT1*, *YAP1*, *HCM1*, *MSN2*, and *SOD2* upon H_2_O_2_ release in *S. cerevisiae* supplemented with resveratrol. For the H_2_O_2_ release quantification, were utilized exponential-growth phase *S. cerevisiae* cultures (O.D._600_ ~ 0.6) grown in SC medium supplemented with 5% glucose. The amplex red hydrogen peroxide assay kit was used for the determination of the H_2_O_2_ release. **a)-f)** Comparison of H_2_O_2_ release between vehicle control and cells supplemented with 100 μM of resveratrol in the WT, *ctt1*Δ, *yap1*Δ, *hcm1*Δ, *msn2*Δ, and *sod2*Δ strains, respectively. The results represent mean values ± standard deviation from three independent experiments. Means were compared with a two-tailed unpaired *t*-test (\**P* < 0.05; \*\**P* < 0.01).

### *Influence of glutathione and resveratrol supplementation in the growth of* ctt1Δ *strain*

The reduced glutathione (GSH) is a well-known antioxidant molecule; we use it to assess whether a pro-oxidant mechanism exerts the detrimental effect of resveratrol upon *ctt1*Δ strain. Exponential growth was utilized as an indicator of resveratrol toxic influence [21]. Resveratrol supplementation decreased the growth of the *ctt1*Δ strain at 5% of glucose (**Fig. 7**). As expected, the supplementation with 100 μM of GSH reverted the negative phenotype exerted by resveratrol upon *ctt1*Δ growth (**Fig. 7**). This result indicates that the negative influence of resveratrol on *ctt1*Δ growth is due to a pro-oxidant effect, which is nullified by the GSH.

**Fig 7.**
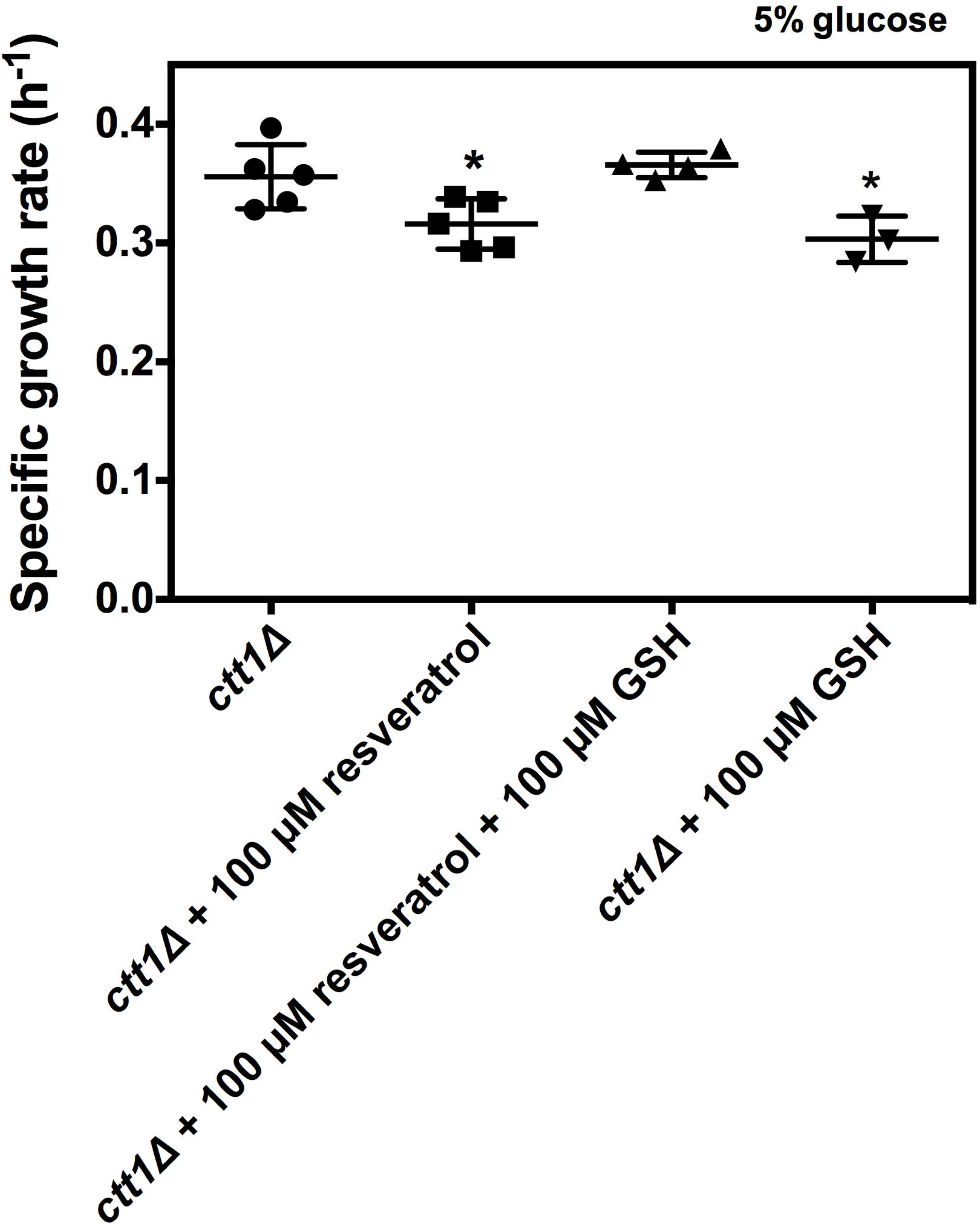
Impact of resveratrol in the exponential growth of *ctt1*Δ strain supplemented with reduced glutathione. The specific growth rate was used as an indicator of the exponential growth of *ctt1*Δ grown in YPD medium at 5% glucose. The results represent mean values ± standard deviation from four to five independent experiments, which include mean values of three technical repetitions. Statistical analyses were performed using one-way ANOVA followed by Dunnett’s test *vs. ctt1*Δ (vehicle control).

### *Effect of resveratrol supplementation in the catalase activity of* S. cerevisiae

The pro-oxidant properties of resveratrol might induce an antioxidant response of cells to counteract the oxidative stress provoked by this phytochemical. For this reason, the *CTT1* gene deletion probably exacerbates the toxic effect of resveratrol. To prove this idea, we evaluate whether resveratrol supplementation increases the catalase activity of *S. cerevisiae*. Unsurprisingly, resveratrol supplementation augmented the catalase activity in comparison with the vehicle control in the WT strain (**Fig. 8a**). Importantly, the *ctt1*Δ strain did not display the increase in the catalase activity induced by resveratrol supplementation (**Fig. 8b**). These data indicate that resveratrol promotes the activation of the catalase activity, possibly by a pro-oxidant mechanism.

**Fig 8.**
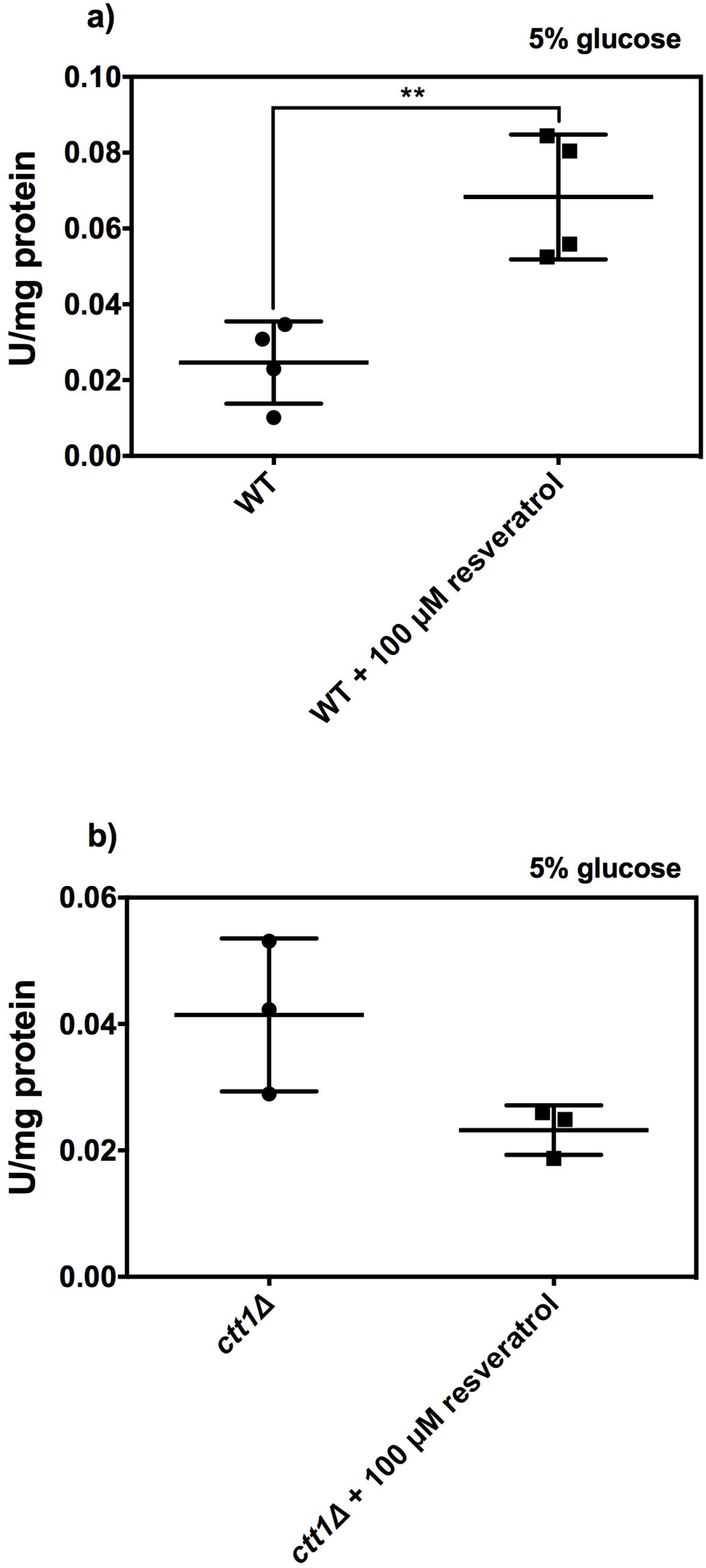
Effect of resveratrol supplementation and *CTT1* gene deletion on catalase activity of *S. cerevisiae.* Catalase activity was measured in exponential-growth phase *S. cerevisiae* cultures (O.D._600_ ~ 0.6) grown in SC medium supplemented with 5% glucose. **a)-b)** Catalase activity comparison between vehicle control and cells supplemented with 100 μM of resveratrol in the WT, and *ctt1* strains, respectively. The results represent mean values ± standard deviation from three to four independent experiments. Means were compared with a two-tailed unpaired *t*-test (\*\**P* < 0.01).

## Discussion

Resveratrol consumption has linked with normalization of the risk factors of some diseases such as colon cancer [17], type 2 diabetes [22], and non-alcoholic fatty liver [23]. Antioxidant activity has been associated with the health benefits showed by resveratrol [24]. However, it is not clear the mechanism by which resveratrol promotes antioxidant activity. The hypothesis that resveratrol inhibits the ETC as its main target, also suggests that resveratrol increases ROS generation produces via reverse electron transport [7]. This idea also explains that cells respond to the oxidative damage caused by resveratrol, inducing their antioxidant systems [5, 7]. The free radical theory of aging postulates that organisms age due to the accumulation of the harmful effects of ROS in cells [14]. For these reasons, we hypothesize that resveratrol shortens CLS of *S. cerevisiae* due to a pro-oxidant activity. Herein, we provide evidence that 100 μM resveratrol supplementation at 5% glucose: 1) shorted the CLS of *CTT1* and *YAP1* genes deleted strains; 2) decreased the H_2_O_2_ release in the WT strain, and maintain unaltered the H_2_O_2_ release in the *ctt1*Δ strain; 3) lessened exponential growth of *ctt1*Δ strain, which was reverted with the adding of GSH; 4) increased catalase activity in the WT strain, a phenotype that was not observed in the *ctt1*Δ strain. Altogether, these results indicate that resveratrol decreases CLS by a pro-oxidant mechanism.

The role that resveratrol exerts on aging is not fully understanding yet. Early studies showed that replicative life span (RLS) was extended with 10, 100, and 500 μM of resveratrol in PSY316AT *S. cerevisiae* cells grown in 2% glucose YPD media [25]. Nonetheless, under the same conditions, it was reported that resveratrol (10 and 100 μM) did not change the RLS of *S. cerevisiae* BY4742 and W303 [26]. Interestingly, it was reported that resveratrol (100 μM) shorted CLS in at 0.5% and 2% of glucose but not at 10% glucose in *S. cerevisiae* BY4742 strain grown in SC medium [8]. A decrease in CLS was also documented with the strain W303-1A grown in minimal medium supplemented with 2% glucose and 100 μM resveratrol [27]. Importantly, CLS lessening was accompanied by an increase of ROS production promoted by resveratrol supplementation [27]. In this study, we found that 100 μM of resveratrol shorted the CLS of the strains deleted in the genes *CTT1* and *YAP1* at 5% glucose (**Fig. 1** **and** **2**). Noteworthy, that *S. cerevisiae* cells grown with 2% glucose or higher concentrations of glucose have a greater ROS production than cells grown at 0.5% glucose [28]. In this sense, the oxidative damage caused by resveratrol supplementation could be enhanced in 5% glucose due to the ROS production provoked at this glucose concentration. In this sense, the deletion of the gene encoding the transcriptional factor Yap1p, required for oxidative stress tolerance and activated by H_2_O_2_, made more sensitive *S. cerevisiae* cells to resveratrol (5 μM) toxicity in YPD and minimal media supplemented with 2% glucose [12]. Besides, supplementation with 5 μM of resveratrol also augmented total ROS levels in *S. cerevisiae* [12]. Interestingly, Yap1p transcriptionally regulates the *CTT1* gene [29], and both gene deletion shorted CLS of *S. cerevisiae* grown with 100 μM resveratrol and 5% glucose (**Fig. 1** **and** **2**). These results pointing out the importance of the cytosolic catalase T to counteract oxidative damage caused by resveratrol, suggesting a possible oxidative-mechanism in the shorten of CLS by this phytochemical.

Several studies have reported the antioxidant response occasioned by resveratrol [30, 31]. Although the molecule has shown the capacity to reduce some oxidant molecules, it is not clear how resveratrol diminishes ROS levels within cells. Even, it has been observed that resveratrol antioxidant response depends on its concentration. For example, supplementation with 5 μM of resveratrol augmented the total ROS levels in *S. cerevisiae* cultures, whereas cultures supplemented with 50 μM of resveratrol decreased it [12]. Besides, also the glucose concentration has an impact on the antioxidant activity of resveratrol. For instance, the H_2_O_2_ release was diminished with 10 μM of resveratrol at 10% glucose, while at 0.5% glucose, the same resveratrol concentration increased the H_2_O_2_ release [8]. We found that the deletion of the *CTT1* gene reverted the diminution of the H_2_O_2_ release by resveratrol supplementation (100 μM) (**Fig. 6**). This result suggests that the H_2_O_2_ release lessening caused by resveratrol is related to a pro-oxidant mechanism countered by antioxidant systems like the cytosolic catalase T. The inhibition of the ETC by resveratrol could explain its pro-oxidant mechanism. Inhibitors of the ETC like resveratrol alter the electron flow rate changing the redox state of some site, lowering respiration rate, and raising ROS production in a given site of the ETC [32]. Resveratrol supplementation disrupts ETC activity between complex I [33] and complex III [21]. In this regard, resveratrol supplementation (30, 50, and 100 μM) increased the basal mitochondrial respiration and decreased the H_2_O_2_ release at 10% of glucose in *S. cerevisiae* cultures [8]. On the contrary, at 0.5% of glucose *S. cerevisiae* cultures displayed a mitochondrial respiration inhibition and an increase in the H_2_O_2_ release with 10 μM of resveratrol [8]. Overall, these data indicate that resveratrol promotes ROS generation via ETC inhibition.

We hypothesize that cells express their antioxidant systems to counteract the oxidative damage caused by resveratrol supplementation. Thus, the antioxidant activity of the cells is responsible for the antioxidant effect showing by resveratrol and not the compound *per se*. Supporting this idea, we found that resveratrol (100 μM) increased catalase activity, and this phenotype was nullified in the *ctt1*Δ strain (**Fig. 8**). Besides, it was also reported that 100 μM resveratrol stimulates catalase and superoxide dismutase activities in *S. cerevisiae* [27]. Additionally, expression of catalase was induced by resveratrol supplementation with 5, 25, 50 and 100 μM in normal human epidermal keratinocytes (NHEK) cells [13]. The pro-oxidant effect of resveratrol has also associated with its toxic influence upon cellular viability. For example, the addition of 25 mM of the antioxidant molecule GSH, augmented the IC_50_ of resveratrol from 247 μM to 747 μM in NHEK cells, from 342.5 μM to 1163 μM in normal human dermal fibroblasts, and from ≥ 150 μM to 445.3 μM in HepG2 liver cells [13]. Likewise, we found that supplementation with 100 μM of GSH rescued the *ctt1*Δ strain of the decrease in exponential growth caused by 100 μM of resveratrol (**Fig. 7**). Altogether, these results indicate that resveratrol pro-oxidative properties play an important role in its toxic effect toward cell viability. Besides, these data suggest that cellular antioxidant systems are responsible for the antioxidant effect showed by resveratrol.

Overall, these results indicate that oxidative stress induced by resveratrol negatively impacts in exponential growth and CLS. Finally, it also suggests that antioxidant effects displayed by resveratrol are due to the cellular antioxidant response.

## Funding

Tecnológico Nacional de México supported LAMP (grant number 5388.19-P).

## Conflict of interest statement

The authors declare that they have no conflict of interest.

## Notes

### Competing Interest Statement

The authors have declared no competing interest.

